# Multi-omics integration of malignant peripheral nerve sheath tumors identifies potential targets based on chromosome 8q status

**DOI:** 10.64898/2026.01.26.701599

**Authors:** Belinda B. Garana, Jiawan Wang, Simge Acar, Gorkem Oztosun, Stavriani Makri, Dana C. Borcherding, Ying Zou, Chelsea Hutchinson-Bunch, Marina A. Gritsenko, Paul D. Piehowski, Christine A. Pratilas, Angela C. Hirbe, Sara J.C. Gosline

**Affiliations:** Pacific Northwest National Laboratory, Richland, Washington, USA; Johns Hopkins University School of Medicine, Baltimore, Maryland, USA; Washington University of St. Louis, St. Louis, Missouri, USA; Oregon Health and Sciences University, Portland, Oregon, USA

**Keywords:** MPNST, proteomics, copy number alteration, EGFR, PLK1

## Abstract

**Background:** Chromosome 8q (chr8q) copy number gain is associated with high-grade transformation in malignant peripheral nerve sheath tumors (MPNST), an aggressive soft tissue tumor with poor outcomes in the high-risk and metastatic settings. Although chr8q gain is associated with inferior overall survival in patients with MPNST, standard of care therapies do not currently consider stratification by genomic features, including chr8q status.

**Methods:** We employed a proteogenomic approach to characterize proteomic and transcriptional programs associated with chr8q and nominate drug targets for potential treatment stratification based on chr8q status. We leveraged our growing library of fully characterized MPNST patient-derived xenografts (PDX) and collected LC-MS/MS global and phospho-proteomics measurements for six of these samples. We then integrated these data with transcriptomics and copy number data to identify molecular changes that are correlated with chr8q copy number. We nominated pathways, transcription factors, and kinases that were differentially active in chr8q gain samples and posited that these samples would respond differently to drugs compared to chr8q wildtype samples. We then tested this hypothesis in vitro.

**Results:** Our results suggest that the chr8q gene *MYC* may be a key driver of downstream effects that can be targetable with inhibitors of PLK1. Conversely, EGFR inhibition may be more effective in MYC-diploid MPNSTs than those with MYC gain. These results nominate candidate pathways and drug classes to target tumor heterogeneity in MPNST through the proteogenomic integration and drug sensitivity prediction in distinct tumor subpopulations.

**Conclusions:** We show that integration of multiomics data can identify specific drug therapies to selectively target tumor cells based on chr8q copy number. This not only provides novel avenues for drug nomination going forward but also may be important for stratifying treatment and mitigating resistance in heterogeneous tumors.

## Background

Malignant peripheral nerve sheath tumors (MPNST) are aggressive soft-tissue sarcomas with poor prognosis. Only twenty to fifty percent of patients over twenty years of age will survive five years from diagnosis.[1] MPNST are usually treated with surgery, with or without chemotherapy and/or radiotherapy. As many as seventy percent of patients develop metastases for which systemic treatment is the only option but is rarely able to achieve durable clinical benefit.[2] To date, no effective targeted therapies have been identified in clinical trials for MPNST, and the median progression free survival (PFS) observed in these trials has been less than two months.[3–5]

Loss of neurofibromin (NF1) and consequent hyperactivation of RAS effector pathways is a key genomic feature required for MPNST development, but other genomic events drive malignant transformation and may confer a more aggressive phenotype [1,6,7] To advance our understanding of biologic features induced by the complex genetic landscape of MPNST, and to offer models suitable for drug testing, we and others have developed patient-derived xenograft (PDX) models of NF1-MPNST[1,8–11] in immunodeficient mice. Using these models, our group found that chromosome 8q (chr8q) gain is associated with high-grade transformation in MPNST and inferior overall survival in 262 patient samples with soft-tissue sarcoma.[8] This study characterizes molecular changes that accompany chr8q gain in these PDX models of MPNST to identify novel targeted treatments for this more aggressive subset.

To examine the proteomic differences in these NF1-MPNST models, we collected liquid chromatography tandem mass spectrometry-based (LC-MS/MS) proteomics using six PDX models. Proteomics offers valuable information about proteins which are directly targetable with small molecule therapeutics, and protein activity through post-translational modifications like phosphorylation. By understanding the proteomic changes driving the aggressive phenotype associated with chr8q gain in MPNST, we posited that we would identify actionable targets with existing, approved drugs; the activity of which has not previously been tested in MPNST. We then integrated the proteomics measurements with copy number data derived from whole genome sequencing and RNA sequencing to identify trends in protein and gene expression associated with chr8q copy number. We employed correlation-based analyses to identify several pathways differentially regulated in MPNST PDX based on chr8q gain and potential drivers which may be coordinating these changes. Additionally, we interrogated the publicly available Cancer Cell Line Encyclopedia for 327 adherent cancer cell lines to identify potential drug candidates. By integrating these multi-omics data sets, we nominated drug candidates to uniquely target chr8q gain and diploid tumors, providing future avenues for combination therapy in MPNST.

## Methods

### LC-MS/MS proteomic measurement of MPNST PDX samples

Proteomics sample preparation and data processing followed protocols developed under the CPTAC consortium.[12–14] mzRefinery was used to process all Thermo RAW files to correct for mass calibration errors before spectra were searched with MS-GF + v9881[15–17] to match against the RefSeq human protein sequence database, release version 38 (https://www.ncbi.nlm.nih.gov/datasets/genome/GCF_000001405.26/). Static carbamidomethylation (+57.0215 Da) on Cys residues, TMT modification (+229.1629 Da) on the peptide N terminus and Lys residues, and dynamic oxidation (+15.9949 Da) on Met residues were considered using MS-GF +. For phosphoproteomics data, IMAC enriched fractions were searched as described above with the addition of dynamic phosphorylation (+79.9663 Da) modification on Ser, Thr, or Tyr residues. Top-scoring assignments for phosphorylation site localization were reported after further processing the phospho-proteomics data using the Ascore algorithm.[18,19] False discovery rate (FDR) calculation was performed using a reversed sequence decoy database approach and results were then filtered to a 1% FDR at the unique protein and peptide level. Parsimonious inference was applied to eliminate redundancy in peptide-to-protein mapping.[20] Proteins and phosphosites were also required to be quantified in at least half of samples. After applying these filters, we obtained 9,013 proteins and 29,545 phosphosites detected in at least 50% of the 6 MPNST PDXs with 2 technical replicates each. After normalizing to the pooled reference channel, values were median normalized across samples and features. Proteomics data can be found in Tables S1 and S2 and is uploaded to Synapse at https://synapse.org/chr8_mpnst.

### Characterization of molecular signatures correlated to chr8q status

We ranked molecular features based on the correlation between their relative expression and median chr8q copy number of the biological sample as described in Figure 1A. Copy number data, depicted in Figure 1B, was derived from our previous work[8] and genes were identified as belonging to chr8q using the positional gene set available from the Molecular Signatures Database (MSigDB, https://www.gsea-msigdb.org/gsea/msigdb/index.jsp)[21] accessed using the function msigdbr from the msigdbr R package as shown in Figure 1C[22]. We performed Spearman correlations between median chr8q copy number and normalized expression values for MPNST PDXs using the function cor.test from the stats R package (http://www.R-project.org/),[23] requiring at least 6 samples per correlation. P-values were adjusted using the Benjamini-Hochberg procedure[24] with the function qvalue from the qvalue R package.[25] Molecular features were then considered significantly correlated with chr8q median copy number if their adjusted p-value was less than or equal to a threshold of 0.05 (shaded values in Figure 1C). We then imposed an additional filter requiring correlated features to have a Spearman rank correlation >0.75 or less than -0.75 (dashed line Figure 1C). Out of 69,797 genes from copy number data, 1,630 out of 19,208 genes from the RNA-Seq data, 208 out of 9,013 proteins and 439 out of 28,866 phospho-sites met this criteria (Figure 1D-E).

**Figure 1.**
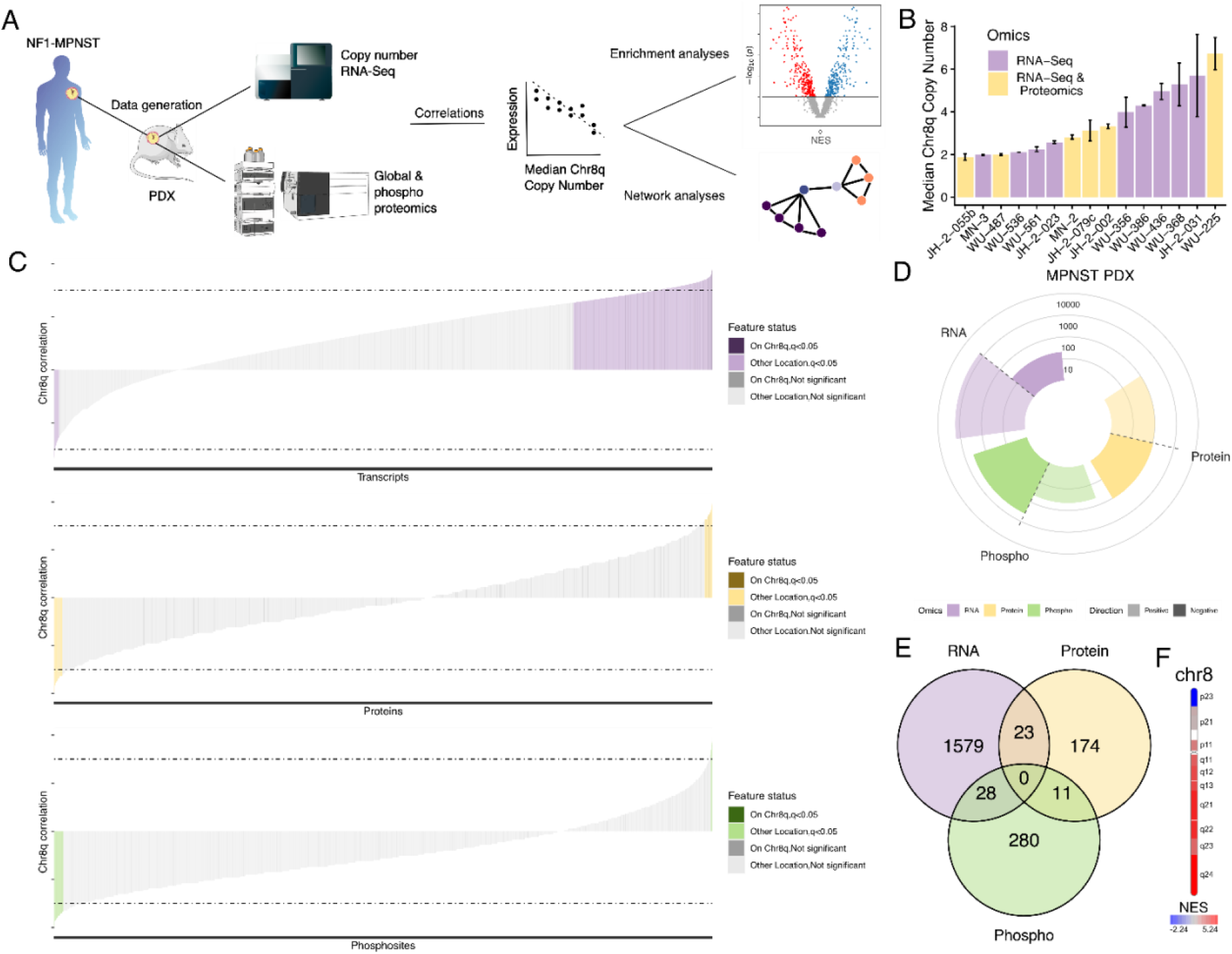
Multi-omics analysis of chr8q gain. (**A**) Overview of our approach. LC-MS proteomics was performed on six of sixteen MPNST PDX samples with RNA-Seq and copy number data with two technical replicates each. Downstream analyses included correlation, enrichment, and network analyses. **(B)** Median chr8q copy number derived from our previous work. Bar colors indicate which data is available for each PDX. (**C**) Correlation of each feature with chr8q copy number. Statistically significant correlations are shaded, while dashed lines indicate the cut-off of 0.75 or -0.75 used to determine correlation. (**D**) Number of features significantly positively or negatively correlated with median chr8q copy number across RNA-Seq (RNA) and LC-MS/MS proteomics (protein) and phospho-proteomics (phospho) levels (adjusted p-value < 0.05). (**E**) Overlap of gene symbols significantly correlated across RNA, protein, and phospho levels. (**F**) Normalized enrichment scores (NES) of chr8 segments based on correlation of copy number with median chr8q copy number.

### Gene set enrichment analysis

To examine global trends in expression correlated with chr8q median copy number, we performed gene set enrichment analysis (GSEA).[26] Specifically, we input Spearman correlation estimates from our molecular signature of chr8q gain (described above) into GSEA and calculated the enrichment of gene sets available in the PID, Hallmark, oncogenic signatures, positional, and Transcription Factor Targets (GTRD) collections on the Molecular Signatures Database (MSigDB, https://www.gsea-msigdb.org/gsea/msigdb/index.jsp)[21] accessed using the function msigdbr from the msigdbr R package.[22] Due to a high percentage of ties in our input rank lists (88-98%; Fig. S1), we permuted tied features (1,000 permutations to generate an ES_ties distribution) in addition to the standard practice of permuting set annotations (another 1,000 permutations to generate an ES_null distribution). We then calculated p-values using the weakest enrichment score (i.e., ES_min) from these tie permutations. We also take a conservative approach for the false discovery rate (FDR) q-values by using the NES_min to identify the fractions of stronger same-signed NES_null and NES_max (i.e., strongest possible scores from other gene sets). Finally, the ES is taken as the mean of the ES_ties distribution and, as standard, the NES is defined as the ES divided by the mean of same-signed ES_null values.

To evaluate the robustness of this adapted GSEA approach which accounts for ties, we: (1) confirmed that chr8q was enriched with chr8q gain as expected (Fig. 1F) and (2) performed a simulation study (Fig. S2). Specifically, we used the 9,013 gene symbols from our global proteomics correlation analysis and generated a representative normal distribution based on that of the real data (normal distribution centered around 0 with standard deviation of 0.5, maximum absolute value of 1, and a median of 90 ties per rank value). Next, we used the MSigDB Hallmark gene set collection of 50 gene sets and added one synthetic gene set consisting of randomly sampled genes from our input list of 9,013. To test the effect of the gene set size on the enrichment, the size of this synthetic gene set was varied from 30 to 300. Once the synthetic gene set was randomly selected, the distribution of the genes in this synthetic gene set was shifted up or down by up to 0.25. This process was repeated a total of 20 times to assess reproducibility. The further this synthetic gene set was shifted, the stronger the enrichment score and more significant the enrichment. However, with no shift (i.e., no difference in distributions between the gene set and the rest of the genes), there were no significant enrichments (i.e., no false discovery), assuring us that this adapted GSEA approach with a distribution is robust even with a highly tied distribution like those of our correlation estimates. We ran the GSEA with simulation on the Hallmarks and transcription factor gene sets and used a p-value cutoff of 0.05 and FDR cutoff of 0.25. 25 of the 29 enriched pathways are depicted in Figure 2A and 25 of the 262 enriched transcription factors in Figure 2B.

**Figure 2.**
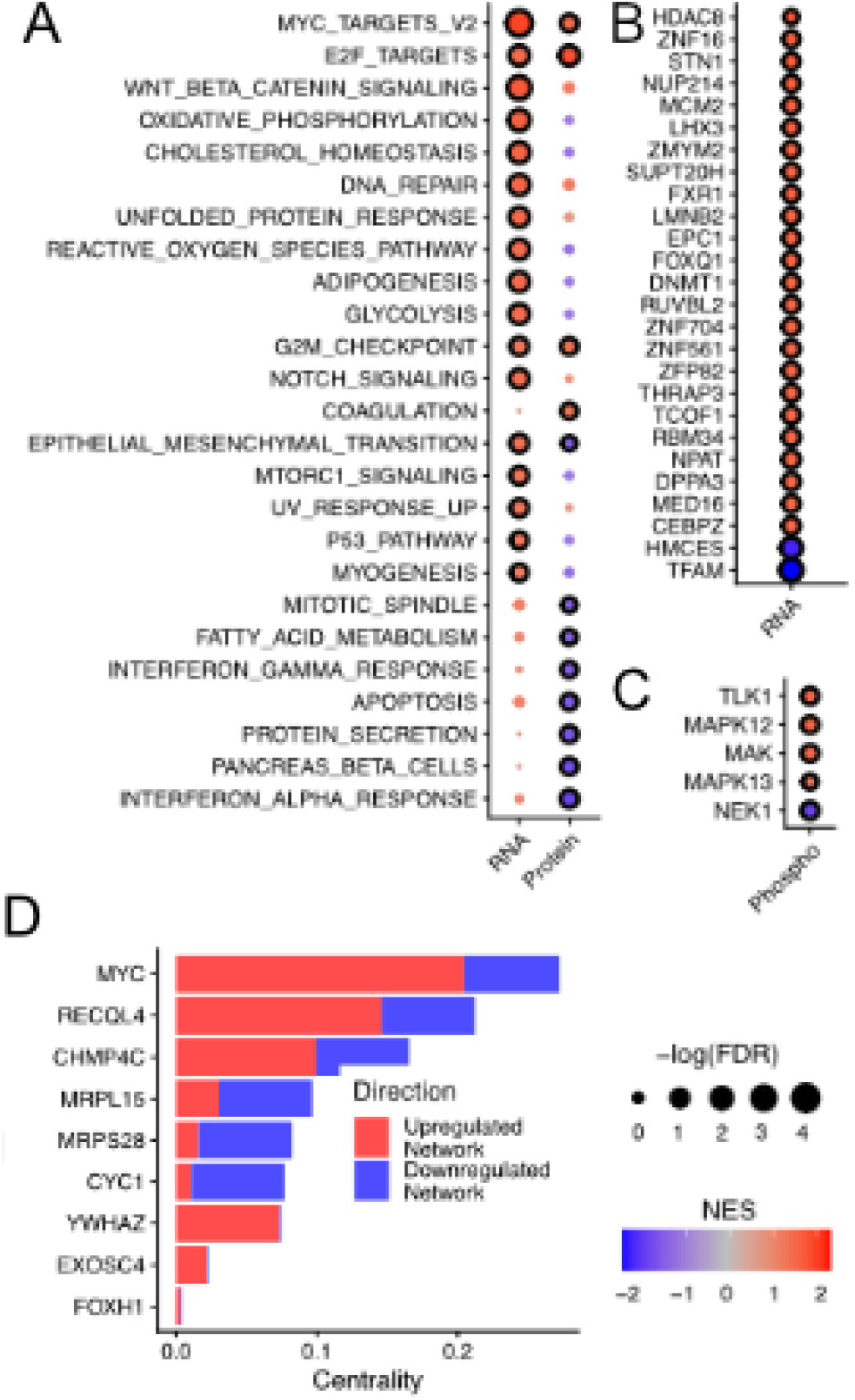
Pathway-level changes caused by chr8q gain measured at transcriptional and post-transcriptional levels. Enrichment based on correlations with median chr8q copy number using either (**A**) Hallmark gene sets from MSigDB across RNA-Seq (RNA) and LC-MS/MS proteomics (protein) levels, (**B**) transcription factor targets from MSigDB for RNA, or (**C**) the kinase-substrate database from PhosphoSite Plus with phosphosite data from LC-MS/MS proteomics. Circle with a border: significant enrichments (p < 0.05 & FDR < 0.25). (**D**) Chr8q gene-associated proteins appearing in both up- and down-regulated networks of transcription factors and kinases ranked by mean centrality.

### Kinase enrichment analysis

To identify kinases which may be differentially active based on chr8 median copy number, we used the kinase-substrate database from PhosphoSitePlus[27] to perform enrichment analysis. The database was downloaded on November 1, 2023. Kinases with at least 6 substrates quantified were evaluated based on enrichment of phospho-site correlations with median chr8q copy number. Kinases with negative enrichments are predicted to be less active with higher chr8q copy number. We used a significance filter of p < 0.05 and adjusted p-value < 0.25 to identify 5 out of 182 (3%) of kinases as enriched per the recommendation for enrichment analysis as shown in Figure 2C.[26]

### Network analysis

To identify potential protein relationships between the features that were correlated with copy number, we built networks from positively and negatively correlated features using the STRING database[28] of physical protein interactions by inputting protein-level features (transcription factors, proteins, and kinases) positively and negatively associated with chr8q copy number, respectively. This approach acts as reinforcement for the enrichment statistics, which can vary across runs due to the stochastic nature of the NES calculation. For the transcription factor and kinase analysis highlighted in Figure 2, this included 260 transcription factors and four kinases (MAPK13, MAK, TLK1, and MAPK12) which were predicted to be more active with higher chr8q copy number or two transcription factors (TFAM and HMCES) and one kinase (NEK1) which were predicted to be less active with higher chr8q copy number. For Figure 3, 101 proteins with expression significantly positively or 107 negatively correlated with median chr8q copy number were added to either the positive or negative networks assembled for Figure 2, respectively. Each network was assembled by querying the STRING database for proteins known to physically interact with these inputs. The full networks consisted of 9,441 positive nodes with 67,958 edges (7,525 nodes with 47,284 edges without protein correlations) or 6,330 negative nodes with 26,034 edges (401 nodes with 812 edges without correlated proteins). Networks were assembled using igraph[29] and then visualized in Cytoscape[30] using RCy3 as depicted in Figure 2D-E.[31] Centrality was calculated using the eigen_centrality function available in the igraph R package.

**Figure 3.**
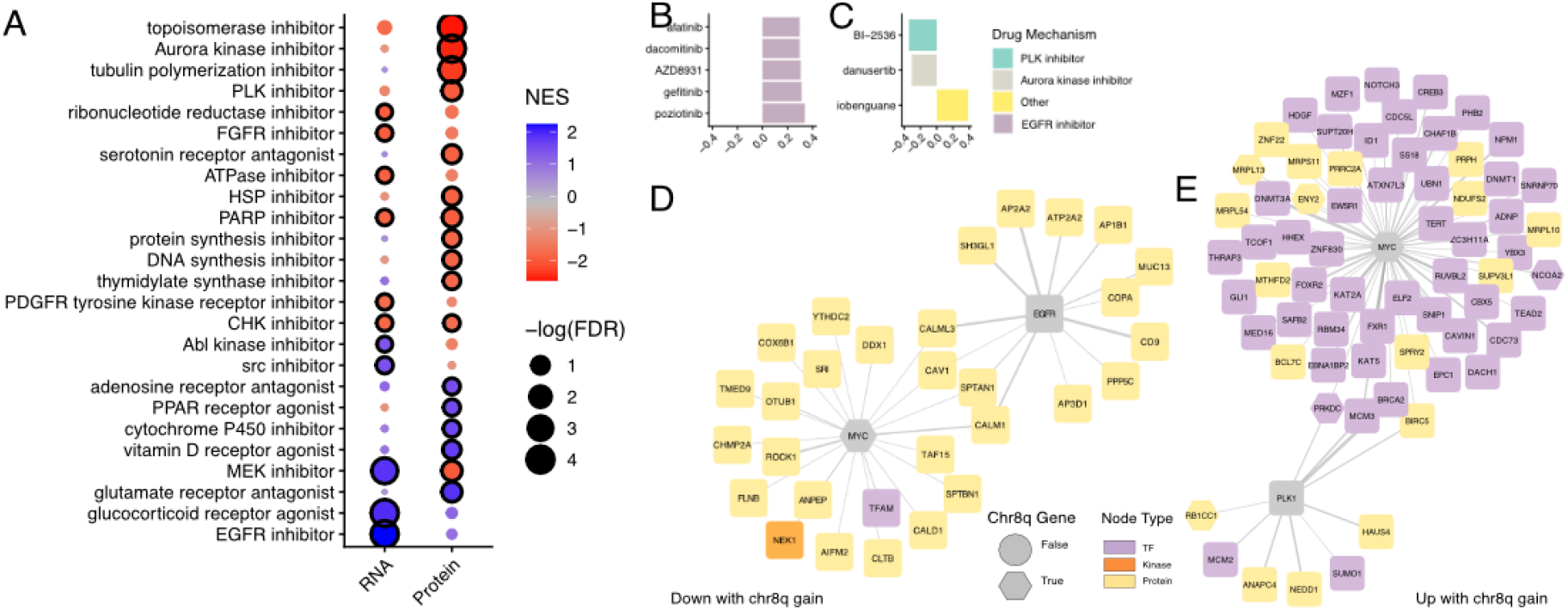
Drug sensitivity analysis reveals potential targets based on chr8q status. (**A**) Enriched drug mechanism sets where NES < 0 indicate potential selective toxicity to MPNSTs with chr8q gain and vice-versa. FDR: false discovery rate. NES: normalized enrichment score. Circled with a border: significant enrichments (p < 0.05 & FDR < 0.25). (**B**) Top five significantly correlated drugs based on absolute Pearson correlation estimates using RNA. (C) Top significantly correlated drugs based on protein data. (**D**) Physical interactions found in STRING database between EGFR, MYC, and proteins up-regulated with higher median chr8q copy number, (E) Physical interactions found in STRING database between PLK1, MYC, and proteins down-regulated with higher median chr8q copy number

### Drug mechanism enrichment analysis

To identify drug mechanisms of action which may be selectively toxic to MPNSTs based on chr8q copy number, we performed drug mechanism enrichment analysis (DMEA).[32] In particular, we input Spearman correlation estimates from our molecular signature of chr8q gain (described above) and queried Cancer Cell Line Encyclopedia (CCLE) global proteomics[33] and RNA-Seq[34] datasets such that the cancer cell lines were ranked based on similarity to our input proteomics and transcriptomics signatures of chr8q gain, respectively. Next, we queried the Profiling Relative Inhibition Simultaneously in Mixtures (PRISM) drug screen[35] to rank 1,448 drugs based on Pearson correlations between drug sensitivity scores (Area Under the Curve, AUC) and these similarity scores in up to 327 adherent cancer cell lines. Finally, we evaluated the enrichment of drugs grouped by 83 known mechanisms of action annotations available from the PRISM drug screen database represented in this drug rank list. P-values and FDR q-values were calculated using 1,000 permutations and a minimum of 6 drugs were required per mechanism of action set. Per the recommendations for DMEA[32] and enrichment analysis in general,[26] we used a p-value cutoff of 0.05 and FDR cutoff of 0.25. Significant mechanisms are depicted in Fig. 3A, while individual drugs are in Fig. 3B-C.

### Cell lines and reagents

Four patient-derived NF1-MPNST cell lines (JH-2-002, JH-2-055d, JH-2-079c, and JH-2-103)[36] were generated in our laboratories at Johns Hopkins University (JH; Baltimore, MD, USA) from biospecimens collected during surgical resection from patients with NF1.[9,11] Material was collected under the Institutional Review Board–approved protocol (#J1649). All patients provided written informed consent. STS26T, ST8814 and NF90.8 cell lines were from Dr. Gregory Riggins (JH; Baltimore, MD, USA). S462 cell line was provided by Dr. Peter Houghton (UT Health San Antonio, TX, USA). NF10.1, NF11.1 were provided by Dr. Margaret Wallace (University of Florida; Gainesville, FL, USA). All cell lines used in these experiments were verified by short-tandem repeat (STR) profiling for cell line authentication at Johns Hopkins University Core Facility, tested negative for mycoplasma contamination, and passaged *in vitro* for fewer than three months after resuscitation. All growth media were supplemented with 10% FBS, 2 mmol/L L-glutamine, and 1% penicillin–streptomycin.

Erlotinib, osimertinib, and volasertib were purchased from MedChemExpress (Monmouth Junction, NJ, USA). BI-2536 was obtained from opnMe by Boehringer Ingelheim. Drugs for *in vitro* studies were dissolved in DMSO to yield 10 or 1 mmol/L stock solutions and stored at −20°C.

### Fluoresence in situ hybridization

Fluoresence in situ hybridization (FISH) was performed on nine MPNST cell lines at the Johns Hopkins Cancer Center Cytogenetics Core Facility. 100 interphase nuclei were scored after FISH with a tri-color probe, where green signal corresponded to IGH (located on chr14q32), red corresponded to MYC (located on chr8q24), and aqua corresponded to cep8 (chromosome enumeration probe for the centromere of chr8). Images are depicted in Fig. 4A. Gain was defined as three or more copies in at least 50% of cells, summarized in Fig 4B-C.

**Figure 4.**
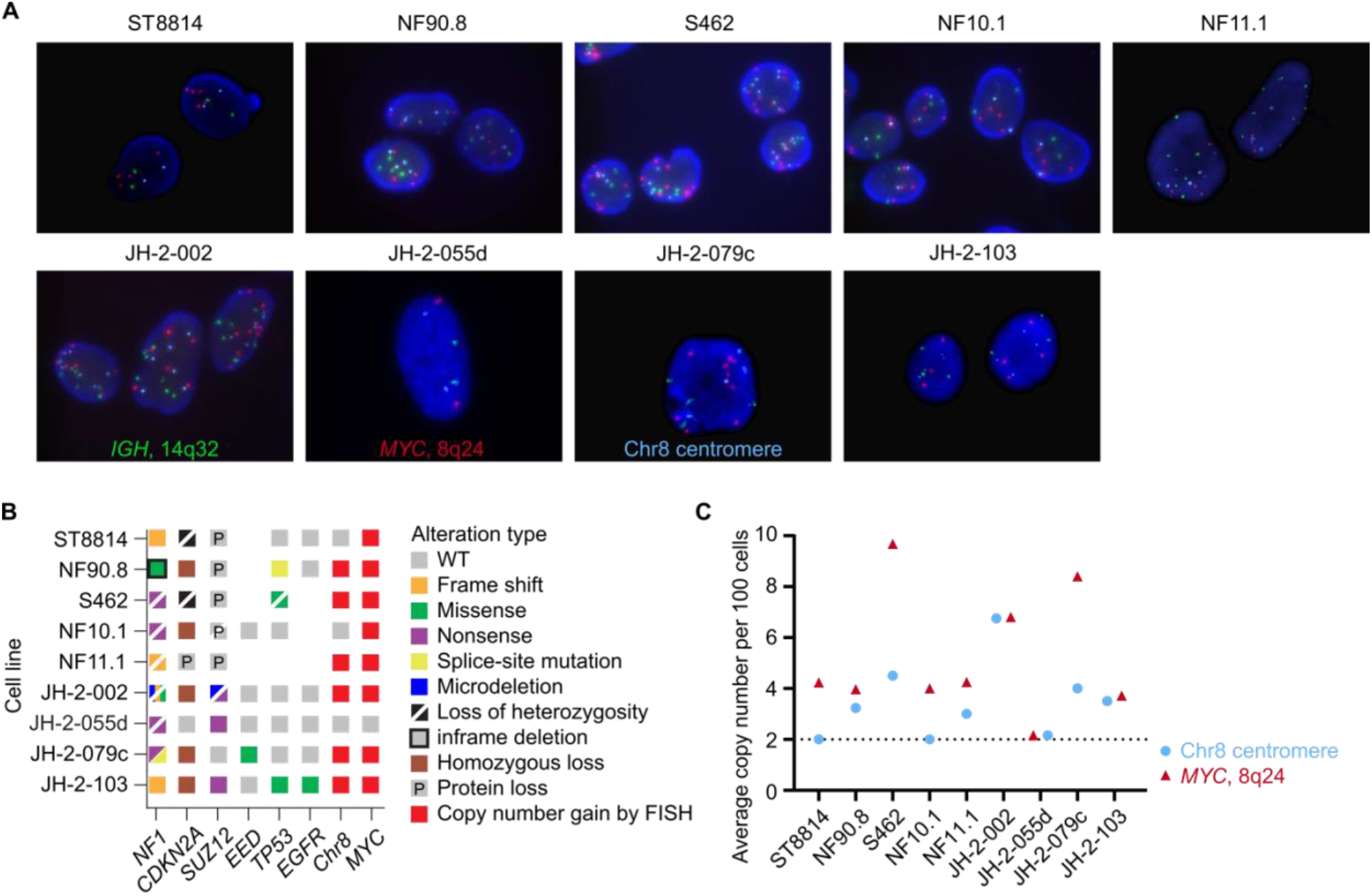
FISH determines chr8 and *MYC* copy number of NF1-MPNST cell lines. (**A**) Representative images of FISH in nine NF1-MPNST cell lines displaying aqua signal for the centromere of chr8, red signal for *MYC*/8q24, and green signal for IGH/14q32. (**B**) OncoPrint integrating key genomic driver features in nine NF1-MPNST cell lines, combining data from our previous study**[36]** with the FISH findings of the current work. (**C**) Average copy numbers of *MYC* or chr8 were quantified per 100 interphase cells in the FISH experiment.

### Cell proliferation assay

JH-2-055d cells were seeded in 96-well plates at 7500 cells per well and other cell lines at 1000-2000 cells per well. A dose range of the compounds indicated was prepared by serial dilutions and then added to the wells containing adherent cells. Cells were incubated with drugs for five days. Cell growth was quantitated using the Cell Counting Kit-8 (Dojindo; Rockville, MD, USA) assay. For each cell line, two to four biological replicates with three to four technical replicates of each condition were measured. Relative survival in the presence of drugs was normalized to the untreated controls after background subtraction and plotted as a function of log_10_ transformed drug doses. The model Sigmoidal dose-response was used to perform the non-linear curve fitting and interpolate the IC50 values using the GraphPad Prism 10. Results and summary statistics are depicted in Fig. 5.

**Figure 5.**
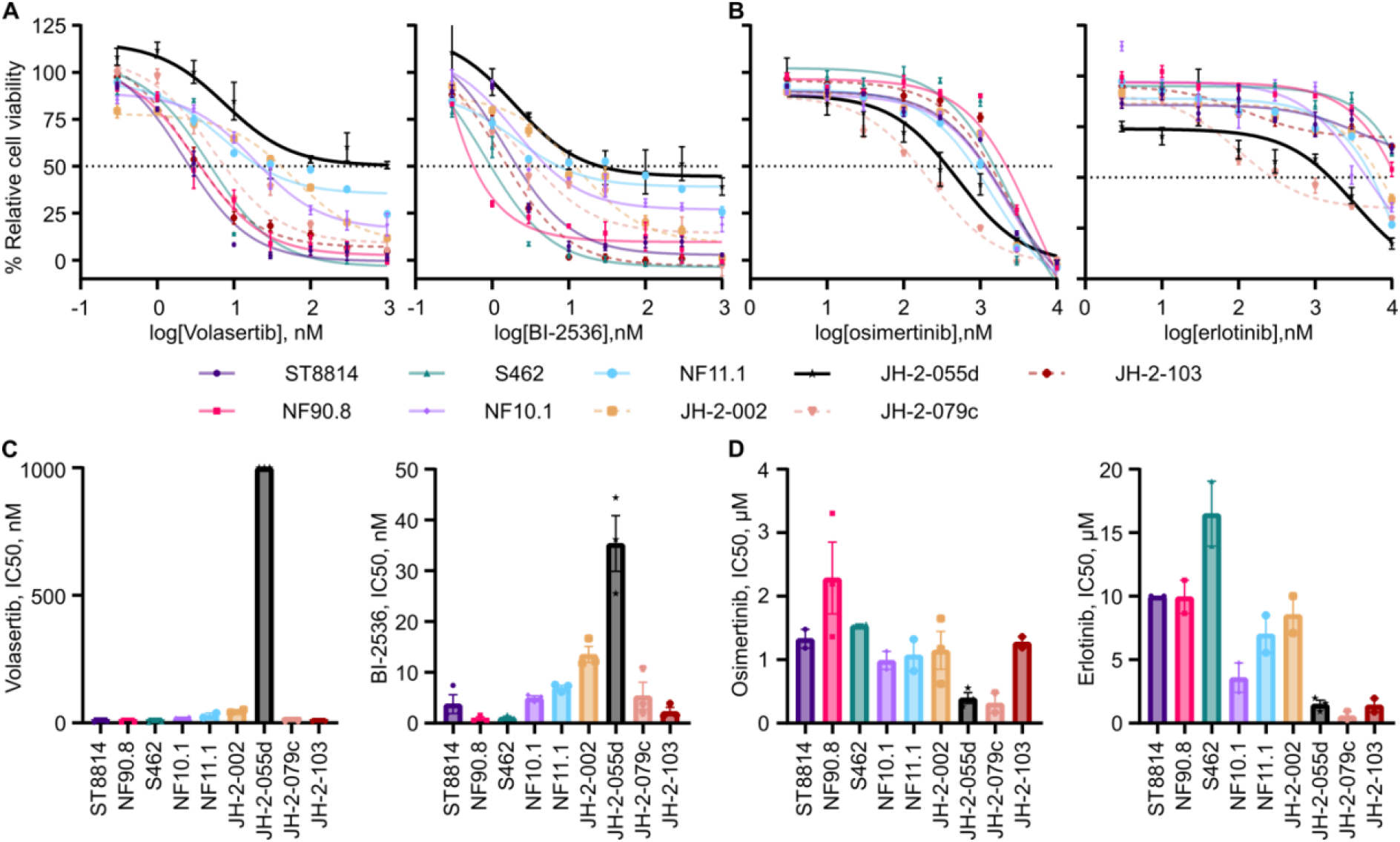
MPNST cell lines have differential response to PLK1 and EGFR inhibitors. (**A-B**) Percent relative cell viability and (**C-D**) IC50 values of MPNST cell lines after 5 days of treatment with either PLK1 inhibitors volasertib or BI-2536 (**A & C**) or EGFR inhibitors erlotinib or osimertinib (**B & D**).

## Results

### Multi-omics analysis illustrates dramatic molecular differences associated with chr8q gain

To identify signaling changes associated with chr8q copy number alteration, we leveraged six previously reported PDX[1,8–11] (Fig. 1A-B) and collected LC-MS/MS measurements of both the global proteome and phosphoproteome using the samples as described in Methods. From these measurements, we identified 9,013 proteins and 29,545 phospho-sites that were present in at least 50% of the samples measured (Table S1-2). To examine the association between chr8q gain and RNA, protein, and phospho-peptide expression, we leveraged existing copy number and RNA-Seq data from these six PDX as well as copy number and RNASeq measurements from other previously published PDX: JH-2-055b, MN-3, WU-536, WU-561, JH-2-023, WU-356, WU-386, WU-436, JH-2-031, and WU-225 [1,8–11]. Next, we performed Spearman rank-based correlations between gene, protein, and phosphosites expression and median chr8q copy number. Here, positive correlations indicate higher expression in chr8q-amplified samples and negative correlations indicate higher expression in chr8q-diploid samples (Fig. 1C, Table S3).

We then used these correlations between chr8q gain and the transcripts, proteins, and phosphosites measured in the same samples to identify systematic patterns across the molecules. We used a Spearman rank correlation statistic and filtered for adjusted p-value < 0.05 and absolute correlation > 0.75 (see Methods). Here we found 1630 (2.19%) transcripts, 208 (2.31%) proteins, and 439 (1.12%) phospho-sites representing 319 proteins to be significantly correlated with chr8q status (Fig. 1D-E; Table S3). Transcripts showed much more statistically significant positive correlation overall (Figure 1C, top panel), consistent with how MYC (located on chr8q24) has been shown to increase overall transcript abundance[38]. Out of the top correlated features based on absolute Spearman correlation estimates across three levels of biological data, we found 103 transcripts, 10 proteins, and 6 phosphosites that are associated with genes that physical reside on chr8q (Fig 1C, darker lines, Table S3), meaning that most associations with chr8q are not simply direct measurements of chr8q gain but instead downstream effects of chr8q. This analysis demonstrates that chr8q gain is associated with nontrivial changes in transcriptomic, proteomic, and phosphoproteomic expression.

We validated this approach by confirming that copy numbers of genes located on chr8q were higher in samples with higher median chr8q copy number by testing for enrichment of chromosome segments in the rankings of our correlations (Fig. 1F). All seven segments of chr8q we quantified were significantly positively enriched (i.e., had greater copy number with higher median chr8q copy number), with chr8q24 (where the oncogene *MYC* is located) being the most strongly enriched (NES = 5.18, p < 1E-3, FDR < 1E-3) (Table S4). One chr8p segment was significantly negatively enriched (chr8p23, NES = -2.2, p < 1E-3, FDR = 1.20E-5) out of five chr8p segments quantified, meaning genes located on these chr8p segments tended to have lower copy number in samples with higher copy numbers of genes located on chr8q[37]. However, three of the five chr8p segments were significantly positively enriched, following the trend of the long arm of chr8 (q). This enrichment of chr8q confirms that our rank-based correlation approach is reflective of changes in line with chr8q copy number.

### Enrichment results show distinct pathway activation patterns with chr8q gain

To quantify pathway-level effects of chr8q gain, we performed gene set enrichment analysis[39] (Table S4 and Fig. 2) using correlations between median chr8q copy number and expression. Positively enriched pathways indicate pathways with higher activity in samples with higher chr8q copy number and negatively enriched gene sets have higher expression in samples with lower chr8q copy number. Our results, depicted in Figures 2A, identify 12 out of 50 Hallmark pathways significantly enriched at the protein level and 17 out of 50 at the RNA level, with 3 pathways in common and significantly positively enriched: MYC targets, E2F targets, and G2M checkpoint (Fig. 2A). Increased activity of MYC with chr8q gain is expected because MYC is located on chr8 itself (specifically chr8q24.21).[40] We also used enrichment analysis to look for protein regulators, including transcription factors using RNA-seq, and kinases using phosphoproteomics (Fig. 2B, Table S4-5). Our analysis suggests 260 transcription factors out of 470 evaluated are significantly more active with higher chr8q copy number but transcription factors TFAM and HMCES may be less active in samples with lower chr8q copy number (Fig. 2B). Additionally, one out of 182 kinases evaluated (NEK1) was predicted to be less active in samples with higher chr8q copy number while four kinases were predicted to be more active (Fig. 2C). To identify which gene(s) on chr8q may be driving these protein regulators, we assembled up- and down-regulated networks of enriched transcription factors and kinases using the STRING physical interaction database (Table S6). We found nine proteins encoded by chr8q genes that were directly interacting with protein regulators and measured the network centrality of these proteins. The results, shown in Fig. 2D, shows that MYC was the most central, particularly in the network of up-regulated features. Metabolic proteins MRPL15 and MRPS28 were most central in networks active in samples with less chr8q amplification, corresponding with the increased apoptosis and metabolism pathways enriched in these same samples in Fig. 2A. Together, these results suggest that chr8q gain drives increased MYC activity across the transcriptional, protein, and phospho-protein levels.

### Drug sensitivity analysis nominates potential targets based on chr8q status

To complement our mechanistic query of potential drug candidates, we employed a data-driven approach that leverages published drug sensitivity data across a panel of 327 adherent cancer cell lines (see Methods, Table S7). Specifically, we ranked each cell line by how similar it was to each our chr8q signatures (derived using transcriptomics or proteomics, see Methods). These rankings were then correlated with drug sensitivity scores (in this case, area under the dose response curve, or AUC), with negative correlations suggesting that MPNSTs with higher chr8q copy number may be more sensitive to the drug of interest, and positive correlations indicating that chr8q-diploid MPNST may be more sensitive. This process leverages large public datasets to identify drugs which may be selectively toxic to MPNSTs based on chr8q copy number.

To score potential drug targets, we grouped the drugs by shared mechanisms of action and performed drug mechanism enrichment analysis (Fig. 3A) to obtain a network enrichment score (NES) for each group of drugs. Drug mechanisms with a negative NES are predicted to be more effective against MPNSTs with higher chr8q copy number whereas those with a positive NES may be more effective against chr8q-diploid MPNSTs. Of the 84 drug mechanisms evaluated 15 were predicted to be more effective in Chr8q gain (negative NES score, Fig. 3A), and 10 were predicted to be more effective with lower amounts of Chr8q gain (positive NES score, Fig. 3A).,Out of 1,448 drugs in the PRISM repurposing database,[35] 83 had sensitivity correlated with expression of protein signatures and 94 were correlated with expression of transcript signatures (adjusted p-value < 0.05). Out of the top 5 correlated drugs based on absolute Pearson correlation, the top drugs predicted to be selective active against chr8q-diploid MPNST are EGFR inhibitors (Fig. 3B) and the top drug predicted to be selectively active against MPNST with chr8q gain include two PLK1 inhibitors and (Fig. 3C). This process allows us to statistically determine which drug mechanisms tend to be selectively effective and also have significantly correlated individual drug candidates.

To explore any potential mechanistic basis for these drug targets, we searched the STRING physical protein interaction network (Fig. 3D-E). Specifically, we sought to define connections between these drug targets, MYC (which we hypothesize may drive the effects of chr8q gain), enriched kinases and transcription factors, and correlated proteins. For the drug target EGFR, we explored connections between MYC, proteins with expression negatively correlated with median chr8q copy number, the one negatively enriched kinase (NEK1), and the negatively enriched transcription factors. We also searched for connections between the drug target PLK1 and MYC, proteins with expression positively correlated with median chr8q copy number, positively enriched kinases, and positively enriched transcription factors. We identified five upregulated proteins and five upregulated transcription factors known to directly interact with PLK1 and thirteen downregulated proteins known to directly interact with EGFR, including three proteins linking PLK1 to MYC and four proteins linking EGFR to MYC. This network analysis shows additional evidence to suggest that PLK1 and EGFR inhibitors might selectively target MPNST in a chr8q copy number-dependent fashion.

### FISH determines chr8 and MYC copy number status across NF1-MPNST cell lines

To validate the differential drug response suggested by our multi-omics analysis, we sought out nine existing NF1-MPNST cell lines to determine if their chr8q status determined their drug response. We first measured chr8/*MYC* copy number status of nine NF1-MPNST cell lines, including the newly established JH-2-055d cell line (corresponding to the same patient from which the PDX JH-2-055b was derived), using fluorescence in situ hybridization (FISH; Tables 1 & S8 and Fig. 4A-C) with probes for the centromere of chromosome 8 (cep8), *MYC* (chr8q24), and *IGH* (chr14q32). Within this nine cell-line panel, six (67%) cell lines exhibited gain in chr8 (3 or more copies of the centromere of chr8), and eight (89%) showed gain in *MYC*/8q24 in over 50% of cells. Only one cell line (JH-2-055d) was *MYC* diploid and three were chr8 diploid (JH-2-055d, NF10.1, and ST8814, Fig. 4C). Given the high prevalence of chr8q gain in MPNST, we are not aware of any other chr8q diploid MPNST cell lines. Specifically, of MPNST cell lines with chr8 gain, most (5 out of 6) had gain in 100% of cells except NF90.8 which had chr8 gain in 84% of cells. Chr8 diploid JH-2-055d revealed chr8 gain in just 11% of cells. Similarly, *MYC* gain occurred in 100% of cells for most cell lines (7 out of 9) tested except for NF90.8 which had *MYC* gain in 98% of cells and *MYC* diploid JH-2-055d where again just 11% of cells had *MYC* gain. For NF11.1, there was non-specific binding observed for the cep8 probe so it could not be distinguished whether NF11.1 had either 3 or 6 copies of centromere 8 but 100% of NF11.1 cells still met the criteria for chr8 gain with at least 3 copies of chr8 (Fig. 4A). We integrated the FISH copy number analysis of chr8 and *MYC* in the current study and molecular alterations in other key genomic drivers of MPNST from our previous report[36] (Fig. 4B), which showed that chr8q gain alongside loss of *NF1, CDKN2A* and PRC2 components *SUZ12, EED*, are common genomic events in MPNST. Notably, the chr8q arm gain was observed more frequently than a gain of the entire chr8 centromere region on this panel of NF1-MPNST cell lines (Fig. 4C). These results are summarized in Table 1 and were then used to stratify cell lines for investigation of drug response, particularly validation of drug candidates nominated above by proteogenomics coupled with drug mechanism analyses.

**Table 1.** Chr8 and *MYC* copy number determined by FISH.

| Cell line | Chr8 gain in $\geq 50\%$ of | *MYC*/ 8q24 gain in $\geq 50\%$ of |
| --- | --- | --- |
| ST8814 | No | 100% |
| NF90.8 | 84% | 98% |
| S462 | 100% | 100% |
| NF10.1 | No | 100% |
| NF11.1 | 100% | 100% |
| JH-2-002 | 100% | 100% |
| JH-2-055d | No | No |
| JH-2-079c | 100% | 100% |
| JH-2-103 | 100% | 100% |

### PLK and EGFR are potential targets depending on MYC status

Drug sensitivity analysis of the PRISM drug repurposing datasets[35] predicted the preferential activity of the PLK1 inhibitors BI-2536 and volasertib (NMS-1286937) against MPNST with chr8q gain, and the selective activity of the EGFR inhibitors poziotinib and pelitinib against chr8q-diploid MPNST (Fig. 3B-C). Given the lack of efficacy and/or toxicity concerns about these two pan-ErbB inhibitors in many other cancers[41–44], and the implication of altered EGFR signaling in a subset of MPNST,[45–47] we focused our analysis on more specific EGFR inhibitors instead, specifically the two FDA approved EGFR inhibitors erlotinib (1^st^ generation) and osimertinib (3^rd^ generation).

To assess the efficacy of PLK1 and EGFR inhibitors on MPNST, we treated nine NF1-MPNST cell lines with PLK1 inhibitors volasertib (clinical stage) and BI-2536 (preclinical compound) and EGFR inhibitors erlotinib and osimertinib for five days (Table S9 and Fig. 5). We grouped the cell lines in Table 1 by chr8 and *MYC* status and compared the difference in cell viability across increasing doses of drug. First, we found chr8 gain lines (all except JH-2-055d) to be more sensitive to PLK1 inhibitors based on IC50 (Fisher combined p = 1.65E-2; p = 5.85E-2 for BI-2536 and p = 4.00E-2 for volasertib) than those without chr8 gain (Fig. 5A & C). Additionally, we found that our chr8 diploid lines were more sensitive to EGFR inhibitors (Fisher combined p = 2.82E-2; p = 1.13E-1 for erlotinib and p = 3.87E-2 for osimertinib) than chr8 gain based on IC50 values (Fig. 5B &D).

Given our hypothesis that the effects of chr8q gain are mainly driven by MYC, we repeated these statistical tests based on the gain status of *MYC* alone instead of all of chr8. Using this approach, we found the difference in IC50 was greater based on *MYC* status than chr8 status: *MYC* gain lines were more sensitive to PLK inhibitors (Fisher combined p = 8.98E-29; p = 1.40E-2 for BI-2536 and p = 9.18E-29 for volasertib; Fig. 5A & C) whereas our *MYC* diploid line was more sensitive to EGFR inhibitors (Fisher combined p = 3.99E-7; p = 3.15E-4 for erlotinib and p = 6.79E-5 for osimertinib; Fig. 5B & D). These *in vitro* data suggests that PLK1 and EGFR may be targetable vulnerabilities in MPNST depending on *MYC* status.

## Discussion

Here we demonstrate that chr8q gain is observed at various levels in MPNST patient-derived xenografts and is associated with changes in levels of RNA and protein expression as well as phosphorylation. Specifically, we used our established cohort of MPNST PDX and associated copy number data to identify genes and proteins that are correlated with chr8q gain (Fig. 1). We also identified pathway level changes including increased MYC activity with chr8q gain, differential activity of transcription factors and kinases, and several potential drug targets (Fig. 2). We used these data to identify putative drug targets to selectively target gain vs. diploid MPNST populations (Fig. 3) and validated these across a suite of genomically diverse cell lines (Fig. 4-5).

Not unexpectedly, MYC was identified as the prominent driver mediating signaling changes associated with chr8q gain, both at the mRNA and protein levels (Fig. 1). This finding was not surprising, given the fact that *MYC* is located on chr8q itself[40] and is a known oncogene.[48] However, our analysis identified many other genes that are associated with chr8q gain but are not physically located on chr8q, and also found significant depletion of chr8p23 in samples with high chr8q copy number (Fig. 1C) suggesting that chr8p and chr8q may be differentially regulated in MPNST. These findings align with previous studies that show both chr8p deletion and chr8q gain are associated with tumor proliferation.[49]

More importantly, our multi-omics analysis was able to identify PLK and EGFR inhibitors to be selectively effective against MPNSTs that are chr8q gain or diploid, respectively. This result was consistent across the cell line correlation data (Fig. 3A-B) as well as network analysis (Fig. 3C-D). Our proteogenomic data analysis revealed that chr8q gain MPNST tumors display significant upregulation of cell proliferation and cell cycle progression, evidenced by the positive enrichment of MYC targets, E2F targets and G2 M checkpoint pathways, at both the RNA and protein levels (Fig. 2A). PLK1 functions downstream of RTK/MEK/ERK signaling, as the master orchestrator of the mitosis phase of cell cycle and has emerged as a promising target for MPNST.[50–52] PLK1 inhibitors have been found to inhibit MYC activity[53] and we have now shown that they demonstrate selectivity for *MYC* gain MPNST (Fig. 5A,C). In the absence of *MYC* gain, chr8q diploid MPNST may be forced to rely heavily on upstream RTK/MEK/ERK pathways for cell survival and proliferation. The ErbB receptor family constitutes a major RTK pathway and specifically, EGFR has been found to promote MPNST transformation via phosphorylation of STAT3.[47] The differential response to PLK1 and EGFR inhibitors observed in these MPNSTs illustrates the importance of considering chr8q status in future MPNST studies.

One limitation of this study that is unavoidable in rare tumors is sample size. With a low sample size, one outlier sample can have a strong effect on correlation estimates. We attempted to mitigate this concern by using Spearman rank-based estimates and permuting tied features for enrichment analyses, though this still caused variation, necessitating network analysis to confirm top hits. We also only had one MPNST PDX in our proteomics study (WU-225) with TP53 loss of function, which is notable since phosphorylation of TP53BP1 contributed to the NEK1 kinase enrichment result (Fig. S3). For our viability assay to test the selective efficacy of PLK1 and EGFR inhibitors, we also only had one *MYC*-diploid MPNST cell line (JH-2-055d) to compare against eight *MYC*-gain lines. This limitation is also difficult to avoid given that our previous study demonstrated that 80% of MPNST harbor highly recurrent chr8q gain.[8] Though our computational analysis included 1,448 drugs spanning 83 drug mechanisms of action screened in 327 adherent cancer cell lines, our drug sensitivity predictions were restricted to these drugs already tested in pan-cancer cell lines, none of which were labeled as MPNST cell lines. Additionally, cell line behavior may differ from that of the patient tumor from which the cell line was derived.

## Conclusions

Overall, this study serves as a multi-omics resource on MPNST patient-derived xenografts, and uses it to identify dramatic changes in expression, pathway activity, and drug sensitivity based on chr8q status. This is a crucial step towards stratifying drug treatments for each MPNST patient based on *MYC* copy number. PLK1 and EGFR inhibitors merit further evaluation in *MYC*-gain and -diploid MPNSTs, respectively. Further interrogating the selectivity of these inhibitors for MPNSTs based on *MYC* status and testing PLK1 and EGFR inhibitors in combination will also be important for targeting tumor heterogeneity, considering both chr8q-gain and -diploid cancer cells may be found in any given tumor.

## Data availability

Data is available upon request on Synapse at https://synapse.org/chr8_mpnst. Processing scripts are publicly available on GitHub at https://github.com/PNNL-CompBio/MPNST_chr8/.

## Acknowledgements

This work was funded by the United States Department of Defense Office of the Congressionally Directed Medical Research Programs (CDMRP) Neurofibromatosis Research Program (NFRP) under W81XWH-22-1-0324 awarded to ACH and HT9425-23-1-0253 awarded to ACH, SJCG, DAL, CAP, and DKW. The mass spectrometry work was performed at the Environmental Molecular Sciences Laboratory, a U.S. Department of Energy National Scientific User Facility located at the Pacific Northwest National Laboratory operated under contract DE-AC05-76RL01830.

## Author information

### Contributions

A.C.H. and S.J.C.G. conceived of the study. S.A. and G.O. generated proteomics samples. C.H., M.A.G., and P.D.P. generated the proteomics data. B.B.G. processed the proteomics data, integrated and analyzed the multi-omics data sets, and led the writing of the manuscript. S.J.C.G. formatted the analysis code for publication and re-use. J.W. performed cell viability experiments and preprocessed samples for FISH. Y.Z. performed analysis on FISH. All authors advised on the study and reviewed the manuscript.

## Supplementary information

Please see separate file: https://wustl.box.com/s/3sowoojeqmc4p8afpcr2x0sj3o5ga8xb

