## Supplementary figures and images for "Multi-omics integration of malignant peripheral nerve sheath tumors identifies potential targets based on chromosome 8q status"

### Fig S1

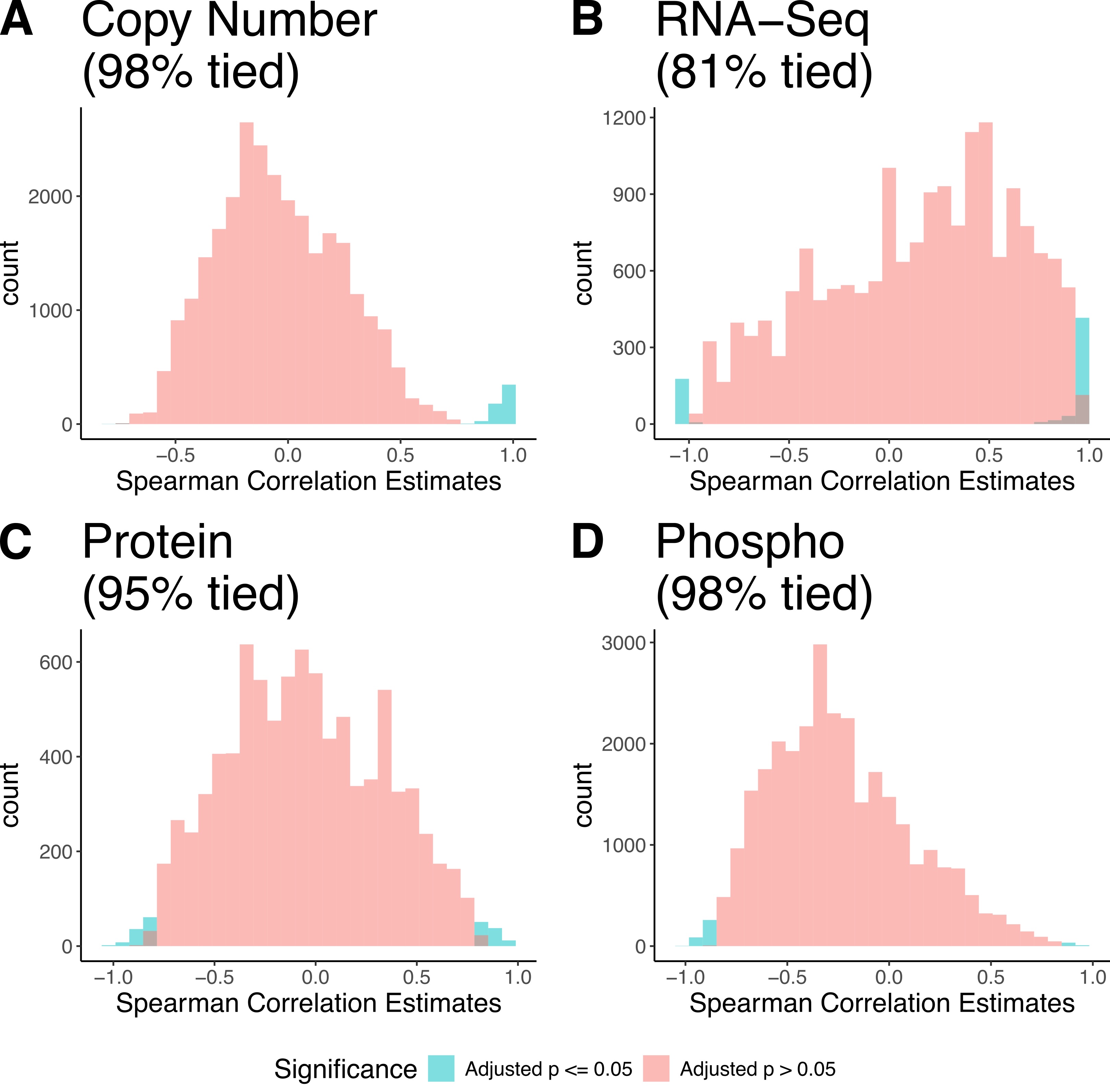

### Fig S2

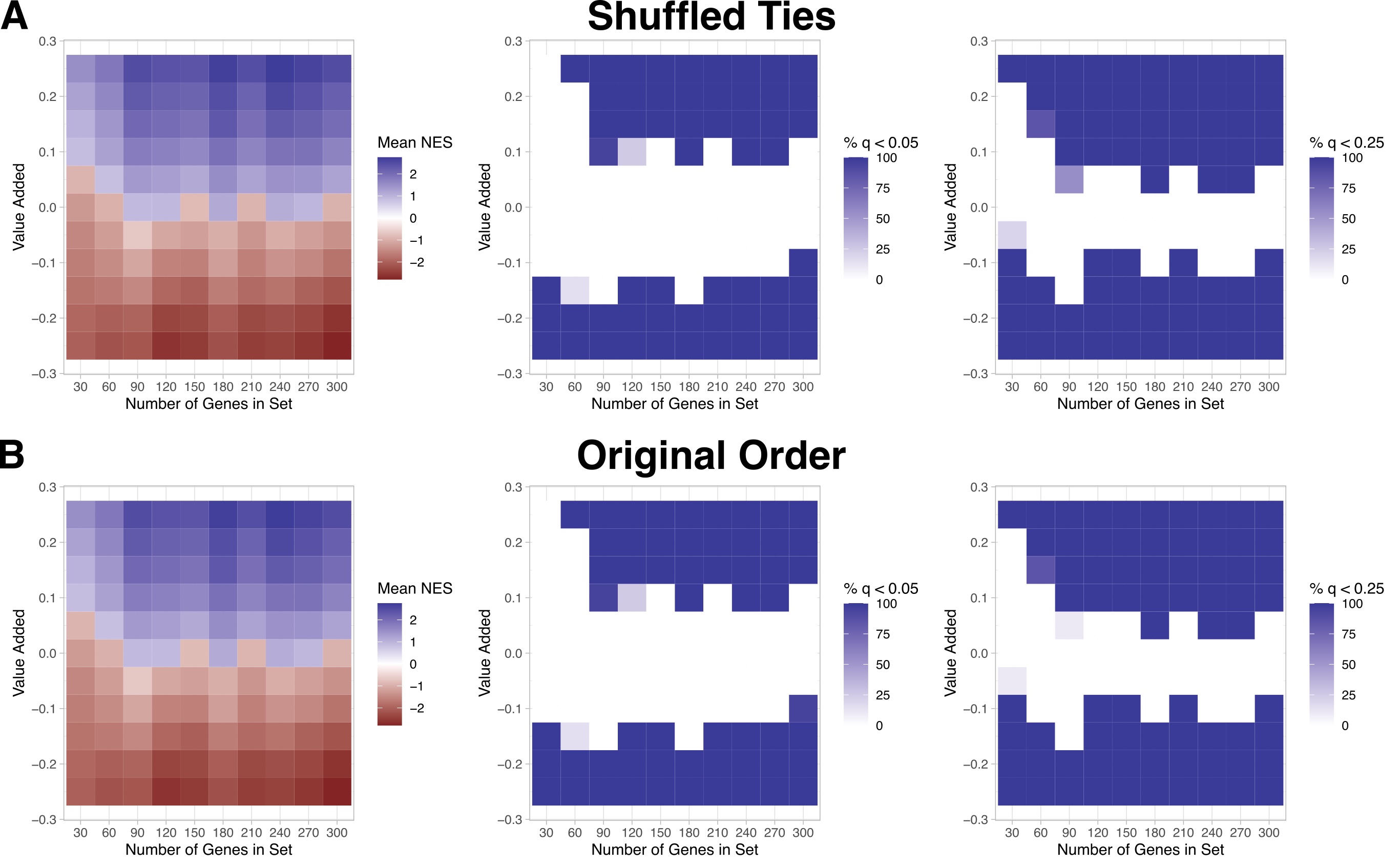

### Fig S3

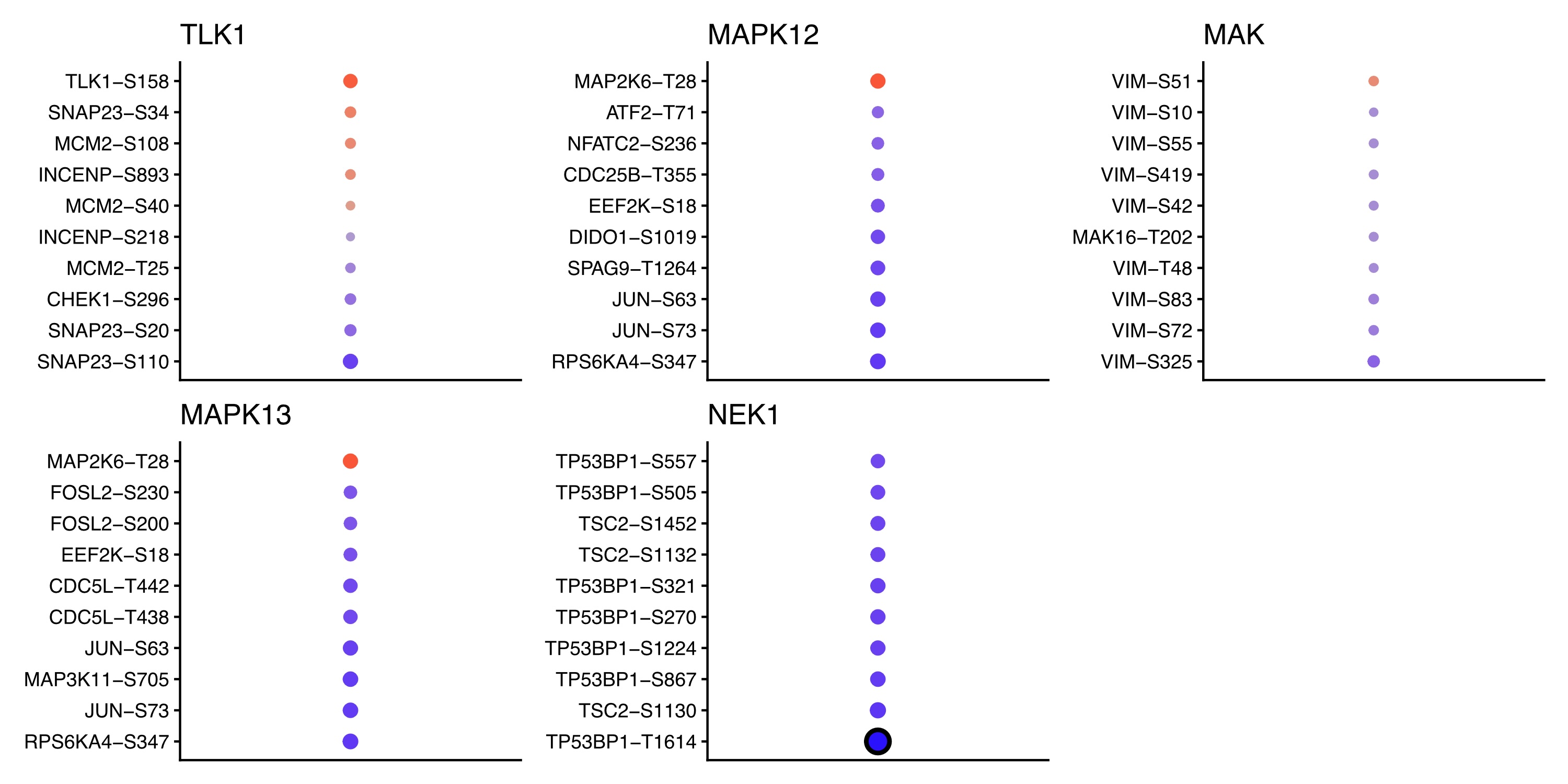
